# The cognitive cost of reducing relapse to cocaine-seeking with mGlu5 allosteric modulators

**DOI:** 10.1101/628388

**Authors:** Christina Gobin, Marek Schwendt

## Abstract

**Rationale:** Cocaine use disorder (CUD) remains difficult to treat with no FDA-approved medications to reduce relapse. Antagonism of metabotropic glutamate receptor 5 (mGlu5) has been demonstrated to decrease cocaine seeking but may also further compromise cognitive function in long-term cocaine users.

**Objectives:** Here we assessed the effect of repeated administration of negative or positive allosteric modulators (NAM or PAM) of mGlu5 on both cognitive performance and (context+cue)-primed cocaine seeking after prolonged abstinence.

**Methods:** Adult male Sprague-Dawley rats underwent 6 days of short-access (1 h/day) and 12 days of long-access (6 h/day) cocaine self-administration. Rats were then trained and tested in a delayed-match-to-sample (DMS) task to establish baseline working memory performance over a five-day block of testing. Next, rats received daily systemic administration of the mGlu5 NAM MTEP (3 mg/kg), mGlu5 PAM CDPPB (30 mg/kg) or vehicle prior to DMS testing during a block of five days, followed by a 5-day washout DMS testing block.

**Results:** MTEP and CDPPB decreased drug seeking in response to cocaine-associated cues after prolonged abstinence. However, repeated treatment with MTEP impaired working memory, while CDPPB had no effects on performance.

**Conclusions:** These results emphasize the relevance of evaluating cognitive function within the context of investigating pharmacotherapies to treat CUD. Further research is needed to determine how two mechanistically different pharmacological compounds can exert the same behavioral effects to reduce cocaine seeking.

## Introduction

Cocaine use disorder (CUD) is a chronic condition characterized by an inability to limit drug intake despite negative consequences (Koob and Volkow 2016). No currently available FDA-approved medications possess the safety profile or display clinical efficacy to reduce relapse rates in CUD. A challenge to treat CUD has been partially attributed to elevated cocaine craving (Gawin and Kleber 1986) and increased reactivity to cocaine-associated cues (Parvaz et al. 2016) that persists over extended periods of abstinence, a phenomenon also replicated in animals (Grimm et al. 2001). A number of neuroadaptations that may be responsible for elevated cocaine-seeking after a drug-free period have been identified in animal studies. One of the best-characterized and replicated findings is that of glutamate release into the nucleus accumbens (NAc) during the reinstatement of drug-seeking in rats (for review see: Scofield et al. 2016). Though dysregulated glutamate homeostasis at the corticostriatal synapses is thought to be primarily responsible (McFarland et al. 2003), other glutamatergic inputs (such as those from the hippocampus, basolateral amygdala and mediodorsal thalamus) may also contribute to the elevated glutamate release during cocaine craving and relapse (Matzeu, Weiss, & Martin-Fardon, 2015; Pelloux et al., 2018; Rogers & See, 2007). Thus, it has been proposed that pharmacotherapies that limit excessive glutamate release or reduce postsynaptic glutamate binding may promote abstinence (Uys and LaLumiere 2008). Due to potentially severe adverse side effects of ionotropic glutamate receptor antagonists, pharmacological modulation of postsynaptic metabotropic glutamate receptor 5 (mGlu5) has been explored as a viable approach for the development of anti-relapse therapies (Olive 2009). A large number of preclinical studies demonstrate that <u>systemic</u> and <u>intra-accumbens</u> administration of mGlu5 antagonists or negative allosteric modulators (NAMs) selectively limit cocaine reward and reduce cocaine-seeking, broadening the appeal of mGlu5 inhibitors as therapeutics for CUD (Chiamulera et al., 2001; Keck et al., 2013, 2014; Kenny et al., 2005; Knackstedt et al., 2014; Knackstedt & Schwendt, 2016; Kumaresan et al., 2009; Lee et al., 2005; Li et al., 2018; Martin-Fardon et al., 2009; Wang et al., 2013),

Despite this evidence advocating for the use of mGlu5 inhibitors in anti-relapse therapy, there are potential complications that may prevent clinical use of these compounds. As mGlu5 receptors play a central role in certain forms of synaptic plasticity (such as long-term depression), it is not surprising that brain-wide inhibition, or ablation of mGlu5 can produce learning and memory side-effects (for review see: Simonyi et al. 2005; Olive 2010). When considering novel approaches to treatment of CUD, it is critical to evaluate and consider potential cognitive side-effects, since up to 30% of dependent cocaine users and 12% of recreational users <u>already</u> exhibit global neurocognitive impairment (Vonmoos et al. 2013). Specifically, deficits spanning multiple domains such as attention, working memory, and executive function have been documented in subjects diagnosed with CUD (Cunha et al., 2013; Goldstein et al., 2004; Jovanovski, Erb, & Zakzanis, 2005; Kübler, Murphy, & Garavan, 2005; Pace-Schott et al., 2008; Potvin et al., 2014; Vonmoos et al., 2013, 2014). Moreover, cognitive deficits may compromise overall treatment outcomes in CUD, as their severity correlates with increased craving (Vonmoos et al. 2013) and relapse susceptibility (Verdejo-Garcia et al. 2014), reduced impulse control (Albein-Urios et al., 2012), or decreased efficacy of cognitive-behavioral therapy (Aharonovich et al. 2006; Streeter et al. 2008). In recent years, several known cognitive enhancers were evaluated as potential treatments for substance use disorders (for review, see: (Brady et al., 2011; Mahoney, 2018). However, these studies revealed no, or only modest benefit of cognitive enhancers on cocaine use or abstinence rates (Dackis et al., 2005; Morgan et al., 2016; Nuijten et al., 2015), and did not concurrently investigate efficacy of these compounds on both cognitive function and cocaine relapse.

This study aimed to address this knowledge gap by evaluating the effects of mGlu5 NAMs and PAMs in an animal model of CUD that allows for the within-subject evaluation of cocaine-seeking and cognitive performance. To that end, we assessed cocaine-seeking in rats with a history of extended (long) access cocaine self-administration after a period of prolonged abstinence (45 days) that corresponded with a time point during which “incubation of cocaine craving”, or elevated (‘incubated’) responses to cocaine-associated cues have been previously demonstrated in both humans (Parvaz et al. 2016) and rodent models (Freeman et al., 2008; Grimm et al., 2001; Lu et al., 2003). During this period, the effects of repeated treatment with mGlu5 allosteric modulators on working memory performance and cocaine-seeking were evaluated in the delayed-match-to-sample (DMS) task and upon re-exposure to cocaine paired context and cues (relapse test). Using this experimental design, we have recently reported that extended-access cocaine self-administration produces not only robust cocaine-seeking, but also persistent working memory deficits that are related to long-term changes in metabolic activity and mGlu5 expression within the prefrontal cortex (Gobin et al. 2019). Here, we tested the following hypotheses: (1) chronic systemic administration of an mGlu5 NAM will impair working memory (2) chronic systemic treatment with an mGlu5 PAM, will improve working memory and (3) acute administration of the mGlu5 NAM, but not PAM will attenuate cocaine-seeking. We chose to evaluate the mGlu5 NAM 3-((2-Methyl-1,3-thiazol-4-yl)ethynyl)pyridine hydrochloride (MTEP) and the mGlu5 PAM 3-Cyano-N-(1,3-diphenyl-1H-pyrazol-5-yl)benzamide (CDPPB), due to their low toxicity as well as the wealth of scientific data supporting <u>anti-relapse</u> properties of MTEP (Keck et al., 2014; Knackstedt & Schwendt, 2016; Knackstedt et al., 2014; Kumaresan et al., 2009) and <u>pro-cognitive</u> properties of CDPPB (Uslaner et al. 2009; Stefani and Moghaddam 2010; Reichel et al. 2011; Fowler et al. 2013; LaCrosse et al. 2014; Marszalek-Grabska et al. 2018).

## Materials and Methods

### Subjects

Adult male Sprague-Dawley rats (Charles River Laboratories; 275g on arrival; N = 38 were first acclimated to the animal facility prior to any manipulation. They were housed individually, maintained on a 12 h reverse light/dark cycle (lights off at 0700), and given *ad libitum* access to food and water, except as noted below. All animal procedures were approved by the Institutional Animal Care and Use Committee of the University of Florida and were performed in accordance with the Guide for the Care and Use of Laboratory Animals. A timeline of the overall experimental design is depicted in Figure 1 with individual behavioral procedures described below. Ten rats were excluded prior to assignment to receive drug or vehicle. Four of these rats died during the cocaine self-administration phase of the experiment, and six failed to reach the predetermined criterion during DMS training (N = 28). Additionally, one rat was considered an outlier during the relapse test according to a Grubb’s test for outliers and was thus excluded from analysis.

**Figure 1.**
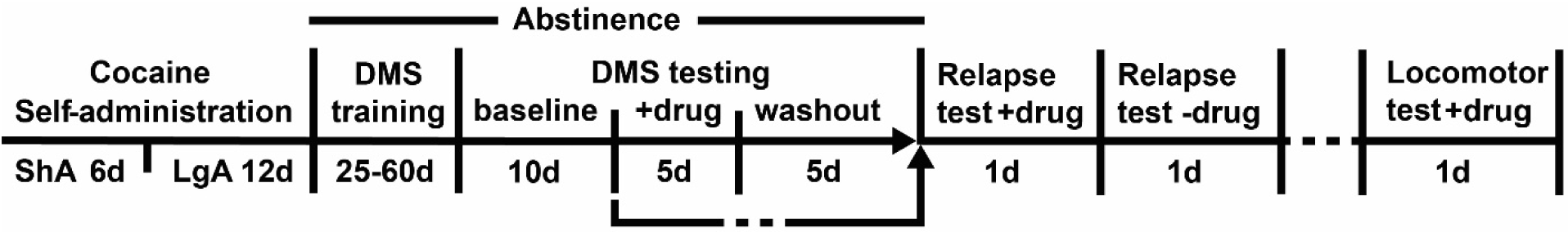
Experimental timeline. DMS – Delayed match-to-sample task. ShA – short, 1 hr access cocaine self-administration. LgA – long, 6 hr access cocaine self-administration.+drug – daily Vehicle, MTEP or CDPPB admininstration, -drug – no drug was administered.

### Drugs

Cocaine hydrochloride was acquired from the NIDA Controlled Substances Program (Research Triangle Institute, NC) and was dissolved in 0.9% sterile physiological saline. MTEP (Abcam Biochemicals, Cambridge, MA) and CDPPB (EAG Laboratories, Maryland Heights, MO) were dissolved in 10% Tween 80 (Sigma-Aldrich) in 0.9% NaCl. Tween 80 (10%) in 0.9% NaCl was used as the vehicle solution. MTEP was neutralized to pH 7.2-7.4 with 1N NaOH. MTEP (3 mg/kg) was administered intraperitoneally (IP) ten minutes prior to behavioral testing. The dose, timing and route of MTEP administration was carefully selected based on previous reports. Specifically, this dose of MTEP resulted in a rapid and full receptor occupancy in adult Sprague-Dawley rats that is maintained for at least one hour (Anderson et al. 2003); reduced cocaine seeking (Martin-Fardon et al. 2009; Hao et al. 2010; Martin-Fardon and Weiss 2012), and was devoid of nonspecific effects on appetitive reinforcement or motor function (Gass et al., 2009; Martin-Fardon et al., 2009). CDPPB was administered subcutaneously (SC) at a dose of 30mg/kg, 20 minutes prior to behavioral testing based on previously published evidence showing pro-cognitive effects of this dose on different types of learning (Stefani and Moghaddam 2010; Olive 2010; Reichel et al. 2011; Fowler et al. 2013) without locomotor side-effects (Gass and Olive 2009).

### Surgery and cocaine self-administration

All rats were implanted with jugular catheters and submitted to cocaine self-administration (SA) as previously described (Gobin and Schwendt 2017; Gobin et al. 2019). Rats were anesthetized with ketamine (87.5 mg/kg, i.p.) and xylazine (5 mg/kg, i.p), and the jugular vein was implanted with Silastic tubing (Dow Corning, Midland, MI). Rats were treated with ketorolac (3 mg/kg, i.p.) to provide analgesia for the first three days of a five-day recovery period. Catheters were flushed daily with 0.1 ml of 100U/ml of heparinized saline and catheter patency was periodically tested by intravenous administration of methohexital sodium (10 mg/mL). Beginning during cocaine SA and lasting throughout the duration of the experiment, rats were restricted to 15-20 g of food per day to maintain ~ 85% of free-feeding weight (Gobin and Schwendt 2017; Gobin et al. 2019).

Rats underwent SA procedures using modified rat operant chambers (30 × 24 × 30 cm; Med Associates, St. Albans, VT) equipped with two standard rat nose poke ports. The chambers were modified to include a mesh wire floor, striped wall pattern, and vanilla scent in order to provide a context that was distinct from the one used in the operant cognitive tasks as described below. Nose poking in the active port resulted in delivery of cocaine (0.35 mg/100 µl infusions) on a FR1 schedule of reinforcement. A 5s presentation of a light + tone cue was paired with each infusion followed by a 20s ‘time-out’ period in which nose-poking was recorded but not reinforced. Nose poking in the inactive port was recorded but did not result in drug or cues. Rats underwent daily 1 hr (short access, ShA) sessions for 6 days, followed by daily 6 hr (long access, LgA) sessions for 12 days. Next, rats entered a prolonged abstinence period during which they began training in the operant DMS task three days following the last day of SA.

### Operant delayed match-to-sample task

Rats were trained and tested in the delayed match-to-sample (DMS) task as previously described (Sloan et al. 2006; Beas et al. 2013; Gobin et al. 2019). Daily 40-minute sessions were conducted in standard rat operant chambers (30 × 24 × 30 cm; Med Associates, St. Albans, VT) equipped with two levers. Within each session, individual trials were composed of three phases - a sample phase, a delay period and a choice phase. During the sample phase, the presentation of the left or right lever was randomly selected by the program such that each lever was presented approximately the same number of times throughout each session. Upon a lever press, the lever was retracted, one sucrose pellet was delivered, and the delay interval was initiated with randomized delay durations. During the choice phase, both levers were extended, and pressing the same lever presented in the sample phase was scored as a correct response and resulted in delivery of one sucrose pellet. Pressing the incorrect lever resulted in a 6s time-out period wherein no sucrose pellet was delivered, the house light was extinguished, and both levers were retracted. Rats were initially trained without any delays between the sample and choice phase. A correction procedure was employed to prevent the development of side biases prior to being trained at two delay sets: short delay set {0s, 1s, 2s, 3s, 4s, 5s, 6s} and intermediate delay set {0s, 2s, 4s, 8s, 12s, 16s}. Rats were required to reach a criterion of ≥ 80% correct for two consecutive days at each training delay set. After reaching criteria, they entered the testing phase for two blocks of five days at the final delay set {0 s, 2s, 4s, 8s, 12s, 18s, 24s}. This phase is designated here as the ‘Baseline’ responding. Next, rats received systemic injections of drug (Vehicle, CDPPB, MTEP) daily prior to DMS testing, for a block of five consecutive days. This was followed by another five-day block of DMS testing, during which no drugs were administered (‘Washout’). One group of rats (1xCDPPB group; n = 7) did not receive any drug treatments; instead, after they completed the baseline DMS testing, they proceeded directly to relapse testing as described below. See the experimental timeline (Fig. 1) for details.

### Relapse tests

On the day after the final day of DMS testing, all rats underwent a 45-min context+cue relapse test in the SA chamber, wherein the cues previously paired with cocaine were presented upon nose poking in the previously active port without cocaine delivery. Rats who received drug treatments prior to DMS testing were administered the same compounds prior to the relapse test. A second context+cue relapse test without prior drug treatments was conducted the following day to assess washout drug effects. Additionally, to assess the effect of a single administration of CDPPB on ‘relapse’ without previous repeated treatment with the compound, one group of rats (n = 7) did not receive drug treatments during DMS testing and instead received a single injection of CDPPB 20 mins prior to the relapse test (Fig.1). A group receiving a single injection of MTEP was not included since it has been previously demonstrated that a single systemic administration of MTEP reduces relapse (Knackstedt & Schwendt, 2016). Since rats reached criterion and finished testing in the DMS test at different time points, the range for the day of relapse testing was 45-80 days. It is unlikely this factor significantly influenced our data, as responding to drug associated cues remains high, but stable between 30 and 100 days of post-cocaine abstinence (Freeman et al. 2008).

### Locomotor Testing

Locomotion was tested in a chamber containing sensors (San Diego Instruments; 40 cm length × 44 cm width × 37 cm) to detect beam breaks in 5-minute bins over the course of one hour. Before locomotor testing, rats received the same drug as previously administered prior to DMS testing and/or relapse testing.

### Statistical analysis

IBM SPSS (Version 25) and GraphPad Prism (Version 5.03) software were used to analyze data. The alpha level was set at 0.05 for all statistical analyses used. A repeated-measures ANOVA was used to analyze discrimination between the active and inactive nose poke ports across the 18 daily sessions of cocaine SA, with Port as the between-subjects factor and Day as the within-subjects factor. To assess cocaine escalation, a repeated-measures ANOVA was conducted to compare mean cocaine intake across the 12 daily LgA sessions of cocaine SA. DMS task acquisition was calculated as the total number of days to meet criterion prior to entering the DMS testing phase. A one-way ANOVA was conducted to assess group differences in number of days to reach criterion prior to DMS testing. During the DMS testing phase, ‘percent correct’ was the primary dependent measure. Percent correct at each delay interval {0 s, 2s, 4s, 8s, 12s, 18s, 24s} was averaged over each five-day testing block. A repeated measures ANOVA was used to assess the effect of treatment on working memory performance for each condition with Block (Baseline, Drug, Washout) as the between-subjects factor and Delay as the within-subject factors. During the relapse tests, our measure of cocaine seeking was operationalized as the number of responses in the previously active port compared to the inactive port. A two-way ANOVA was used to assess group differences in the number of nose pokes in the active and inactive ports during both context+cue relapse test days. A one-way ANOVA was conducted to assess group differences in locomotor performance. Bonferroni post-hoc tests were used when appropriate.

## Results

### Rats escalated cocaine intake during long-access cocaine self-administration

All rats demonstrated successful discrimination between the active and inactive nose poke port throughout SA (Fig. 2A). A repeated-measures ANOVA revealed main effects of Port (*F*(1, 62) = 279.2, *p* < 0.0001) and Day (*F*(17, 1054) = 39.45, *p* < 0.0001) with a Port × Day interaction (*F*(17, 1054) = 45.32, *p* < 0.0001). A repeated measures ANOVA with Greenhouse-Geisser correction showed that rats displayed an escalation of cocaine intake (mg/kg) over the 12 days of LgA sessions (*F*(5.20, 171.68) = 15.69, *p* < 0.001). Post-hoc tests using Bonferroni correction revealed that cocaine intake on LgA days 4-12 were significantly higher than on day 1 (*p* < 0.05; Fig. 2B). On average, total cocaine intake (± SEM) for the whole period of self-administration was 821.05 ± 32.86 mg/kg, with the long-access self-administration phase accounting for the majority of cocaine intake, 778.04 ± 29.75 mg/kg.

**Figure 2.**
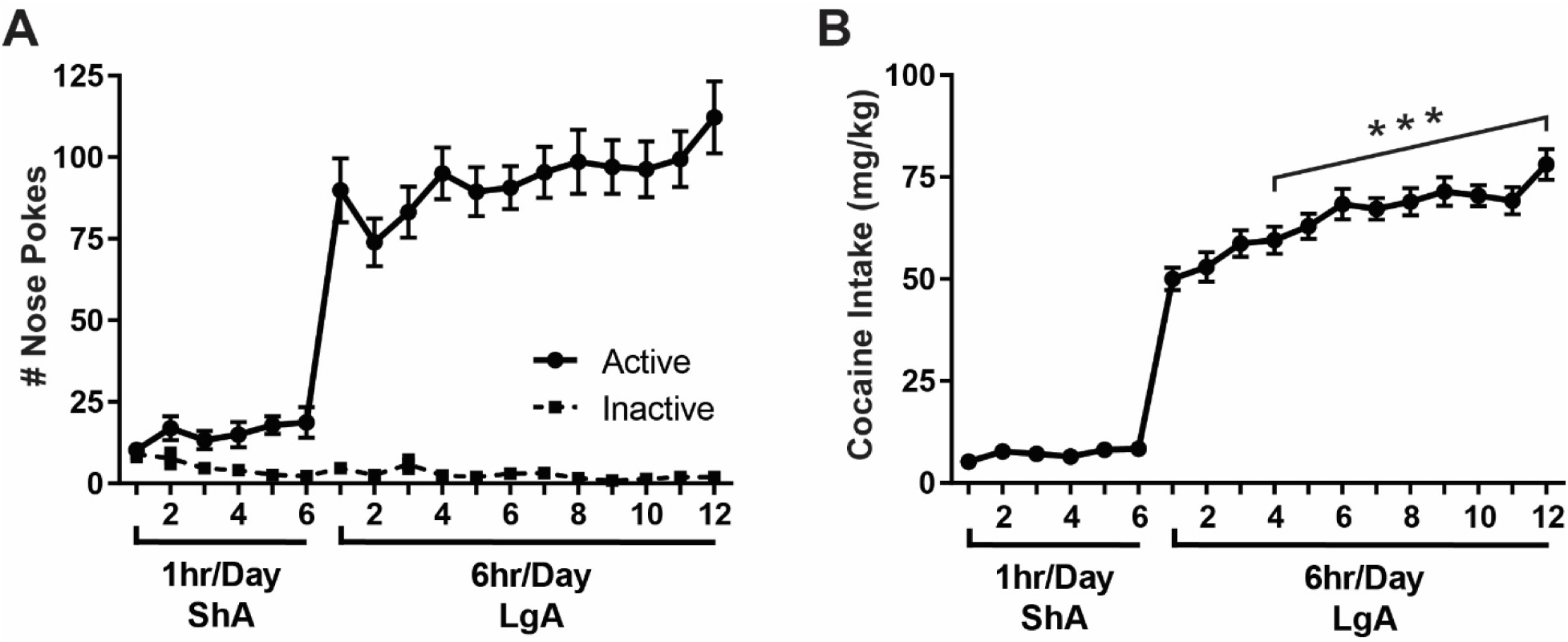
Cocaine self-administration. A) Rats discriminated between nose poking in the active versus inactive port throughout the self-administration. There were main effects of Port and Day as well as a Port × Day interaction. B) Rats showed escalation of cocaine intake on days 4-12 of LgA cocaine self-admininstration. ***p < 0.001 vs. intake on Day 1 of LgA cocaine self-administration, N = 28.

### Rats did not differ in DMS task acquisition or baseline performance prior to treatment

A one-way ANOVA did not reveal a difference between groups on task acquisition prior to treatment (*F*(3, 25) = 1.35, n.s; Fig. 3A). Prior to behavioral pharmacology testing, rats were matched to treatment groups such that there were no group differences in total cocaine intake (*F*(3, 25) = 2.76, n.s.) and baseline DMS performance across all delays (no effect of Group *F*(3,25) = 1.58, n.s.; and no interaction *F*(18, 144) = 1.21, n.s).

**Figure 3.**
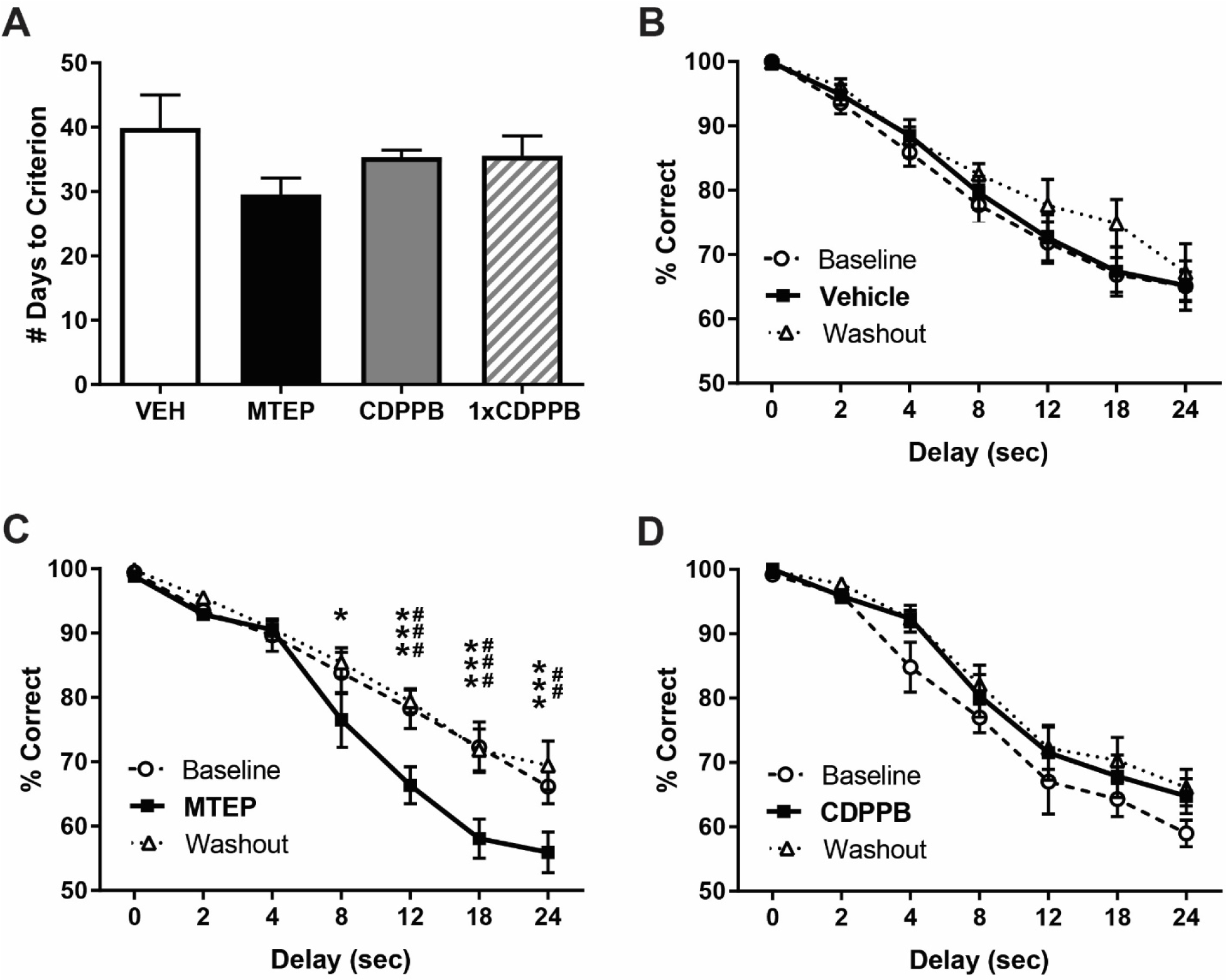
Delayed math-to-sample task. A) Animals assigned to four treatment groups did not display pre-exisitng differences in task acquisition (number of days to reach criterion). B) Treatment with vehicle did not affect task performance (% correct) compared to baseline and washout blocks. C) Task performance significantly decreased during the MTEP (3 mg/kg, i.p.) treatment block compared to the baseline block at 12s, 18s, and 24s and compared to the washout block at 8s,12s, 18s, and 24s. D) Treatment with CDPPB (30 mg/kg, i.p.) did not alter task performance compared to baseline or washout blocks. ##p < 0.01, ###p < 0.001 MTEP vs. Baseline, *p < 0.05, ***p<0.001 MTEP vs. Washout. n = 6-8/group.

### MTEP impaired while CDPPB had neutral effects on working memory performance

For the Vehicle treatment group, main effects of Delay (*F*(6, 98) = 29.27, *p* < 0.0001) and Block (*F*(2, 98) = 9.55, *p* < 0.001) were found without a significant Delay × Block interaction (Fig. 3B). For the MTEP treatment group, main effects of Delay (*F*(6, 84) = 51.52, *p* < 0.0001) and Block (*F*(2, 84) = 26.99, *p* < 0.0001) as well as a Delay × Block interaction (*F*(12, 84) = 3.05, *p* < 0.01) were found (Fig. 3C). Bonferroni post-tests revealed that MTEP treatment reduced the percent correct at the 12s, 18s, and 24s delays compared to baseline (*ps* < 0.01-0.001). MTEP performance was also impaired at the 8s, 12s, 18s, and 24s compared to the washout block (*ps* < 0.001-0.05). (Fig. 3C). For the CDPPB treatment group, main effects of Delay (*F*(6, 70) = 50.43, *p* < 0.0001) and Block (*F*(2, 70) = 9.06, *p* < 0.001) were found without a significant Delay × Block interaction (Fig. 3D). The effects of MTEP on working memory are not likely due to decreased appetitive behavior, as groups did not differ in total number of trials completed averaged over the five-day treatment block (*F*(2, 20) = 3.38, n.s). The mean (± SEM) number of trials completed over the 5-day treatment period was: 102 ± 4.98 (Vehicle group), 101.7 ± 3.55 (MTEP group), and 116.5 ± 1.12 (CDPPB group).

### Both MTEP and CDPPB decreased drug-seeking in a context+cue relapse test

A two-way ANOVA revealed main effects of Port (*F*(1, 23) = 41.75, *p* < 0.0001), Treatment (*F*(3, 23) = 3.76, *p* < 0.05) and a Port × Treatment interaction (*F*(3, 23) = 3.85, *p* < 0.05) during the first relapse test (Relapse test+drug; Fig. 4A). Bonferroni post-hoc tests showed that rats in the MTEP, repeated CDPPB, and 1xCDPPB groups exhibited less nose pokes in the previously active port compared to the Vehicle treatment group (*ps* < 0.01-0.001). Only rats in the Vehicle group significantly differed in the average number of nose pokes in the active port versus the inactive port (*p* < 0.001). On the following day (Relapse test - drug), a two-way ANOVA revealed main effects of Port (F(1, 23) = 58.77, p < 0.0001), but no effect of Treatment (F(3, 23) = 1.36, n.s) and no significant Port × Treatment interaction (F(3, 23) = 2.17, n.s).

**Figure 4.**
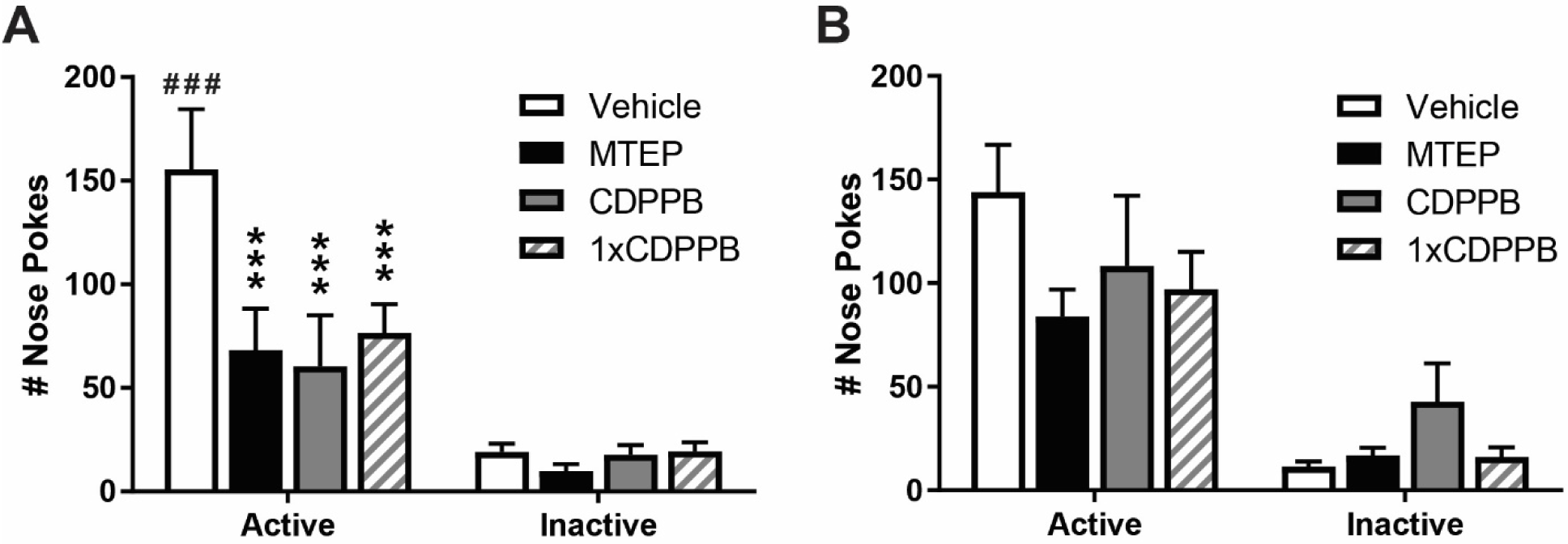
Relapse to cocaine-seeking. A) Rats with a history of vehicle prior to DMS testing and relapse exhibited greater # nose pokes in the active port compared to rats with a history of MTEP (3 mg/kg, i.p.) and CDPPB (30 mg/kg, i.p.) prior to DMS testing and relapse as well as compared to rats given only a single administration of CDPPB prior to relapse. Vehicle-treated rats also nose poked more in the active nose port compared to the inactive nose port. B) Groups did not differ on the # nose pokes in the previously active port on day 2. ***p<0.001 vs. Veh active port nose pokes. ###p<0.001 vs. inactive port nose pokes within each group. n = 5-8/group.

### Neither drug treatment altered spontaneous novelty-induced locomotion

We evaluated whether the doses of the drugs used in this study affected locomotor activity. A one-way ANOVA showed that treatment groups (Vehicle, MTEP, CDPPB) did not differ in the overall locomotion in a novel environment, measured as average number of beam breaks during a 1 hour session (*F*(2, 21) = 1.74, n.s.). Mean number of beam breaks (± SEM) for each group was as follows: Vehicle (5023 ± 385), MTEP (4370 ± 229) and CDPPB (5529 ± 387).

## Discussion

Our data indicate that both allosteric inhibition and activation of mGlu5 reduces relapse to cocaine-seeking in rats, even after prolonged abstinence. At the same time, the dose of MTEP that decreased relapse also produced significant working memory deficits. On the other hand, allosteric activation of mGlu5 with CDPPB at a dose that reduced relapse, spared working memory, indicating that mGlu5 PAMs might offer safer approach to reduce cocaine relapse.

Expanding on previous studies showing that systemic MTEP reduces cocaine-seeking (Keck et al., 2014; Knackstedt & Schwendt, 2016; Kumaresan et al., 2009), we are the first to report that systemic MTEP administration also attenuates (context+cue)-elicited cocaine-seeking following extended access cocaine SA and prolonged abstinence (45-80 days). In other words, MTEP attenuates cocaine-seeking in animals that typically show sustained elevation of drug-seeking and resistance to extinction, behavior previously described as ‘incubation of cocaine craving’ (Grimm et al. 2001; Lu et al. 2003; Freeman et al. 2008). We hypothesize that systemic MTEP is partially targeting the NAc to control drug seeking as previous studies report that MTEP microinfused into the NAc is sufficient to reproduce the effects of systemic MTEP administration (Knackstedt et al., 2014; Li et al., 2018; Wang et al., 2013). However, mGlu5 receptors are widely distributed in other brain regions (Rodrigues et al., 2002; Romano et al., 1995; Shigemoto et al., 1993) known to play a role in cue-elicited cocaine-seeking (Carelli et al. 2003; Pelloux et al. 2018). As previous studies have shown that intra-vmPFC or intra-dlSTR blockade of mGlu5 do not reduce relapse to cocaine-seeking (Ben-Shahar et al., 2013; Knackstedt et al., 2014), anatomically separate populations of mGlu5 in the brain may play distinct roles in regulating drug-seeking. Thus, possible contribution of mGlu5 receptors outside of the NAc to drug-seeking behavior warrants further, more comprehensive evaluation.

The MTEP-induced disruption of post-cocaine working memory performance was most apparent during longer delays (8-24s), suggesting a delay-dependent (or load-dependent) impairment. These effects of MTEP were acute and did not persist past the drug treatment period (denoted here as washout). We recently found that the same regimen of cocaine self-administration used in the present study produced deficits in working memory performance that persisted over weeks of drug-free abstinence (Gobin et al. 2019). As all rats used in the current study had a cocaine history, their working memory ability was likely already impaired prior to MTEP administration, indicating that treatment with this class of drugs may produce further cognitive impairment in recovering addicts. The deficits produced by MTEP here are of comparable magnitude to those resulting from mPFC lesion in the same task (Sloan et al. 2006). In addition to the mPFC, lesion and inactivation studies using match-to-sample and nonmatch-to-sample operant tasks have identified that circuitry supporting ‘normal’ working memory performance also includes the dmSTR (Akhlaghpour et al. 2016). Thus, we hypothesize that systemic MTEP may be impairing working memory through inhibition of specific cortico-striatal circuits. The acute effects of MTEP here also suggest that mGlu5 activity is critical for ongoing working memory performance. Within the mPFC, mGlu5 is predominantly expressed postsynaptically on neurons projecting to various striatal regions, including the NAc and dmSTR (Romano et al. 1995). Indeed, administration of MTEP directly into the mPFC impairs working memory using the analogous DMS task (Hernandez et al. 2018).

As MTEP impaired working memory, we predicted that a mGlu5 PAM would enhance working memory. Here, we tested the effects of CDPPB, an mGlu5 PAM shown in other tasks to exert pro-cognitive effects (Uslaner et al. 2009; Stefani and Moghaddam 2010; Reichel et al. 2011; LaCrosse et al. 2014). Surprisingly, we did not detect an effect of CDPPB on post-cocaine working memory performance in rats. Previously, the same dose of CDPPB (30mg/kg) improved methamphetamine-induced episodic memory deficits (Reichel et al. 2011), as well as MK-801-induced deficits in cognitive flexibility (Stefani and Moghaddam 2010) and spatial learning (Fowler et al. 2013). Several factors could account for the lack of effect found in our study. In these studies, CDPPB effects were observed in significantly impaired animals, with the discrimination or response accuracy near, or at chance (Stefani and Moghaddam 2010; Reichel et al. 2011). And further, in these tasks facilitatory effects of CDPPB typically applied to new learning. In contrast, we have shown that chronic cocaine produces sustained, but relatively mild cognitive impairment in rats (Gobin and Schwendt 2017; Gobin et al. 2019) and this study). Also in this study, CDPPB was evaluated on rats who had an extensive history of DMS training. And finally, while the difficulty of the task increased during testing (longer delays), new learning was not required to master the DMS task. To assess possible delayed effects of CDPPB on recall of the DMS task rules, we conducted a continuation study (maintaining individual group designations) wherein rats’ working memory performance was re-evaluated after relapse and locomotor tests were completed (~ 6 days after the end of the washout period). Under these conditions, we did observe working memory enhancement in rats previously treated with CDPPB (Suppl. Fig. 1), but not in rats with a history of Vehicle or MTEP treatment (not shown).

In contrast to the divergent effects MTEP and CDPPB on working memory, both drugs reduced cocaine-seeking triggered by cocaine-associated context and cues. The anti-relapse effects of CDPPB were not due to cumulative effects of prior repeated treatment with this drug (5-day CDPPB treatment regimen during DMS testing), as single CDPPB administration (20 mins prior to the relapse test) also reduced cocaine-seeking. As intra-NAc administration of orthosteric mGlu5 agonist CHPG has been shown to promote cocaine-seeking or conditioned effects of cocaine (Wang et al. 2013; Benneyworth et al. 2019), potent anti-relapse effects of CDPPB may seem surprising. However, the relapse test as conducted in the current study is analogous to an extinction session on day 1 in studies that demonstrated extinction-enhancing effects of CDPPB (Olive 2010; Cleva et al. 2011). Importantly, on extinction day 1, these studies show a similar CDPPB-induced decrease in overall responding to drug cues (Cleva et al. 2011). Thus, the CDPPB-induced reduction in drug seeking may be attributed to acute enhancement of extinction learning occurring already within the first session.

Allosteric modulators typically regulate functional receptor activity specifically in brain areas where the endogenous agonist exerts its physiological effects (Stansley and Conn 2019). Thus, it is possible that MTEP and CDPPB (two mechanistically different compounds) may be acting at opposing brain regions to modulate mGlu5 function related to the expression of distinct behaviors (decreased motivation to seek the drug vs. enhancement of extinction learning) that would result in the same outcome (decrease overall responding in the relapse test). MTEP may be decreasing the ability of drug-associated cues to drive PrL brain activation and behavior (Martin-Fardon et al. 2009; Kumaresan et al. 2009; Keck et al. 2014) via an mGlu5-mediated inhibition of neuronal excitability within this region (Timmer and Steketee 2012) or the NA core (Wang et al. 2013), but not infralimbic cortex (IL, (Ben-Shahar et al. 2013), whereas CDPPB may be facilitating within-session extinction learning via an mGlu5-mediated activation within the IL (Ben-Shahar et al. 2013). Furthermore, the efficacy of these modulators in enhancing mGlu5 activity depends on the concentration of endogenous/orthostatic agonist (extracellular glutamate) and on mGlu5 agonist-modulator cooperativity factor (Stansley and Conn 2019). As such, region-specific dysregulation of mGlu5 signaling (Gobin et al. 2019), mGlu5 function (Ben-Shahar et al. 2013) and glutamate homeostasis (Knackstedt & Kalivas, 2009) after chronic cocaine may further determine circuit-specific effects of systemic MTEP and CDPPB administration.

The current study also assessed cocaine-seeking one day after the initial relapse test (Relapse test +drug), to evaluate possible carryover effects of prior pharmacological manipulations on cocaine-seeking (Relapse test -drug). Here, we show that prior treatment with MTEP or CDPPB did not affect drug-seeking in this subsequent test. This finding is in agreement with our previous observations that single intra-dSTR, but not systemic MTEP administration interferes with post-relapse extinction learning (Knackstedt et al. 2014; Knackstedt and Schwendt 2016). It is possible that repeated MTEP administration is needed to decelerate extinction (Kim et al. 2014). In the case of CDPPB, we again predict that its repeated daily administration is necessary to produce extinction-enhancing effects (Olive 2010). It should be noted that one study observed an effect of CDPPB or MTEP administration on cocaine-seeking during a subsequent ‘undrugged’ extinction test (Perry et al. 2016). However, in this study, either compound was administered prior to a CS+ extinction session that was separate from operant extinction. Another factor not addressed in the aforementioned studies, is the resistance to extinction in rats with a history of extended access to cocaine (Grimm et al. 2001; Lu et al. 2003; Freeman et al. 2008), and also observed in our Vehicle-treated rats. Thus, it is possible that longer extinction sessions (45mins vs. 2 hrs) are necessary to observe retarding or faciliatory effects of mGlu5 PAMs on extinction in these animals.

## Conclusions

The current study is the first to investigate the effects of positive and negative mGlu5 allosteric modulation on both cognitive performance and persistent cocaine-seeking. We found that repeated systemic administration of the mGlu5 NAM MTEP impaired working memory performance, while mGlu5 PAM CDPPB had no effect. Interestingly, both compounds attenuated relapse to cocaine-seeking triggered by combined discrete and contextual cocaine cues, without affecting subsequent extinction learning. We hypothesize that systemic administration of MTEP reduces the ability of mGlu5 receptors within the cortico-accumbal pathway to mediate cocaine-seeking, at the cost of inhibiting working memory processes. On the other hand, the ability of CDPPB to exert the same anti-relapse effect (though free of cognitive side-effects) makes mGlu5 allosteric modulation an attractive target for the development of future therapies of CUD. These initial observations motivate further research into the brain circuitry and neurobiological mechanisms through which mGlu5 PAMs reduce drug seeking.

## Acknowledgements

The authors thank Dr. Jennifer Bizon and Dr. Joseph McQuail for their input on the design of the DMS task and Dr. Lori Knackstedt for her valuable comments during manuscript preparation. We also thank Mr. Spencer Berman, Mr. Kyle Fontaine, Ms. Carly Wallace and Ms. Vera Monlux for their assistance with conducting behavioral studies.

## Conflict of interest

The authors declare that they do not have conflicts of interest.

## Funding information

This research has been supported by the University of Florida McKnight Brain Institute pilot award (MS).

## Compliance with ethical standards

All experimental protocols were approved by the University of Florida Institutional Animal Care and Use Committee and consistent with guidelines of the NIH Guide for the Care and Use of Laboratory Animals.

---

**Supplementary Fig. 1.**
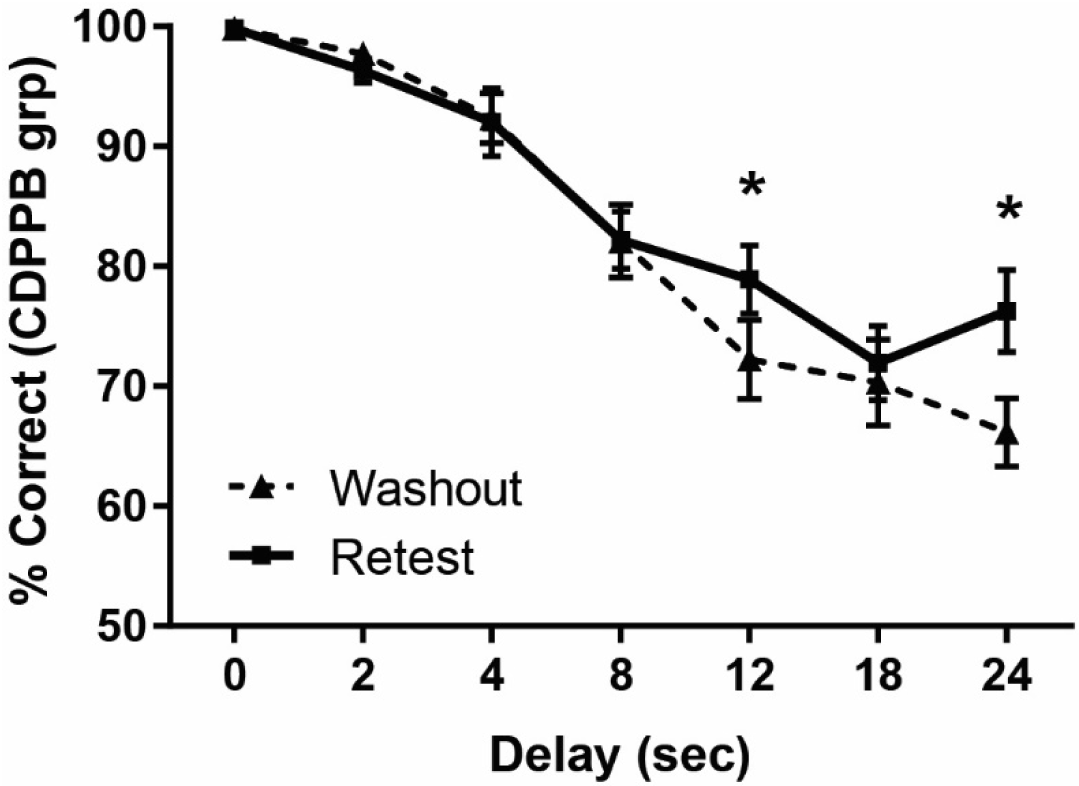
Retesting of CDPPB-treated rats. A) Six days after the end of initial washout period, rats previously treated with CDPPB (this time, no drug was administered) showed an improvement in DMS performance at the 12s and 24s delays, as compared to the past Washout block. Data represent an average of 5 daily DMS sessions. n = 6. *p<0.05 vs. Washout.

## Notes

#### Summary of Updates

Funding information has been updated.

